# Sex-specific associations between education-related genetic factors and fertility extend beyond educational attainment

**DOI:** 10.64898/2026.04.29.721701

**Authors:** Ivan A. Kuznetsov, Alexandros Giannelis, Estonian Biobank Research Team, Kelli Lehto, Triin Laisk, Cornelius A. Rietveld, Uku Vainik, Vasili Pankratov

**Affiliations:** Institute of Genomics, University of Tartu, Tartu, Estonia; Department of Applied Economics, Erasmus School of Economics, Erasmus University Rotterdam, Rotterdam, The Netherlands; Institute of Psychology, University of Tartu, Tartu, Estonia; Montreal Neurological Institute, McGill University, Montreal, Canada; Bioinformatics Research Centre, Aarhus University, Aarhus, Denmark

**Keywords:** Fertility, Educational attainment, Genetic factors, Sex differences, Mechanisms

## Abstract

Educational attainment (EA) is strongly associated with fertility, yet the mechanisms underlying this relationship remain poorly understood. Using data from ∼40,000 women and ∼10,000 men in the Estonian Biobank, we examine sex-specific associations between EA polygenic scores (*PGS_EA_*) and completed fertility. The *PGS_EA_* captures an individual’s genetic predisposition to completing higher levels of education, and can also be constructed separately for cognitive and non-cognitive EA components. We find that *PGS_EA_* is negatively associated with fertility among women, with a significantly stronger association for the non-cognitive than the cognitive EA polygenic score. The association of *PGS_EA_* with fertility is only partly explained by EA and is moderated by EA and reproductive timing, changing sign across age-at-first-pregnancy strata. Within-family estimates are not attenuated relative to population estimates, consistent with a predominant role of direct genetic effects. Among men, associations between *PGS_EA_* and fertility are non-significant or slightly positive. Overall, education-related genetic variation is associated with fertility through pathways largely independent of EA, operating through sex-specific and timing-dependent mechanisms. These results contribute to our understanding of the micro-level mechanisms shaping macro-level demographic change.

## Introduction

The demographic transition – marked by sustained declines in mortality followed by a decline in fertility – has reshaped population age structures and labor markets worldwide. In contemporary low-fertility societies, variation in completed fertility increasingly reflects heterogeneity in socioeconomic attainment, partnership trajectories, and life-course timing (1, 2). Among these factors, educational attainment (EA) occupies a central position. Across a variety of countries, higher levels of education are typically associated with delayed childbearing and lower completed fertility, particularly among women (3–5). This negative educational gradient in fertility has been interpreted through multiple frameworks, including opportunity cost models (6, 7), gender equity theory (8, 9), and ideational change (10).

More recently, advances in molecular genetics have introduced a complementary perspective: genetic variants associated with educational attainment also predict reproductive outcomes. Genome-wide association studies (GWAS) have identified thousands of genetic loci associated with EA, enabling the construction of polygenic scores (PGS; also referred to as “polygenic indices” or PGI) that aggregate these associations into a single measure predictive of EA (11, 12). The same studies have demonstrated robust associations between the EA PGS (*PGS_EA_*) and years of schooling in European-ancestry samples. Subsequent research has extended this work by examining whether EA-associated genetic variants are linked to fertility outcomes, often interpreted through the lens of natural selection in modern populations. Several studies have reported that higher *PGS_EA_* is associated with lower lifetime reproductive success (13–16). Educational attainment also shows a negative genetic correlation with fertility (17). These findings suggest that alleles related to higher educational attainment may be under negative selection in contemporary low-fertility contexts.

Associations between *PGS_EA_* and fertility do not necessarily mirror associations between educational attainment and fertility. For example, while differences in the education–fertility association across groups in the United States have been shown to be substantial, corresponding estimates for *PGS_EA_* have been nearly identical (16). Likewise, in the Icelandic population, *PGS_EA_* has been negatively associated with fertility among men, despite a positive (though not statistically significant) association between educational attainment and fertility. Moreover, even when phenotypic and genetic associations share the same direction, educational attainment may not fully account for the association between *PGS_EA_* and fertility, pointing to alternative mechanisms linking EA-related genetic variation to fertility (13). Thus, analyses based on polygenic scores provide a complementary perspective, allowing the characterization of factors associated with, and potentially causally linked to, fertility.

Several mechanisms may underlie negative associations between education-related genetic loci and fertility, as well as between education level and fertility. First, classical human capital models emphasize the substitution effect: as wages rise with education, the opportunity cost of childbearing increases, incentivizing delayed and reduced fertility (6, 7). Under this framework, genetic variants associated with higher EA may reduce fertility by increasing earnings potential and career investment (14). At the same time, the income effect operates in the opposite direction: higher income can facilitate childrearing by relaxing budget constraints. Given that, in advanced economies, the opportunity costs of childrearing are higher for women (1), we expect substitution effects to be more pronounced among women, whereas income effects may be more salient among men (5, 18–22). Such asymmetry implies potential sex-specific associations between *PGS_EA_* and fertility.

Second, education is linked to shifts in preferences, aspirations, and life-course sequencing. Higher EA is associated with later partnership formation and delayed age at first birth, both of which are associated with completed fertility (5, 23, 24). Genetic predispositions toward non-cognitive traits – such as conscientiousness, future orientation, or risk aversion – may further influence reproductive timing. Recent work has decomposed genetic associations with EA into cognitive and non-cognitive components (25), enabling the construction of polygenic scores for these distinct components (*PGS_Cog_* and *PGS_NonCog_*, respectively). These components have shown opposite genetic correlations with personality traits such as conscientiousness and agreeableness. If non-cognitive traits shape partnership stability or career prioritization, they may exert stronger effects on reproductive outcomes than cognitive ability per se. However, no differences in genetic correlations between cognitive and non-cognitive EA components and number of children have been found in either women or men (25).

Third, associations between PGS and fertility may reflect indirect genetic effects operating through parental socioeconomic status or assortative mating, rather than causal influences of an individual’s genotype (26). Distinguishing direct genetic effects from environmentally induced associations is essential. Within-family designs, such as adjustment for parental PGS (26) or comparisons of siblings differing in their PGS and trait values (27), provide a more stringent test of direct genetic effects by controlling for shared family environment and residual population stratification. Evidence from such designs remains limited, particularly for fertility-related outcomes (28).

Importantly, gene-environment interplay may generate context-dependent associations. Comparative demographic research shows that educational systems, labor market structures, and family policies can amplify, attenuate, or reverse the fertility correlates of education-related traits (29–31). For example, generous parental leave and childcare provision may mitigate the career–family trade-off, weakening negative educational gradients in fertility (32, 33). Correspondingly, the magnitude of associations between EA-related genetic variants and fertility appear to vary also across social and demographic environments within societies (5, 14, 15).

To date, nearly all evidence linking EA-associated alleles to fertility derives from the US or a few Western European cohorts (5, 13–17, 34, 35). This narrow empirical base substantially limits inference about the universality and generalizability of observed patterns. The documented heterogeneity in both EA–fertility and *PGS_EA_*–fertility associations underscores the importance of examining genetically informed fertility gradients in underrepresented populations undergoing distinct social and institutional transformations.

Estonia provides a particularly informative context for studying genetically informed fertility patterns. Fertility in Estonia fell below replacement already in the 1920s and, unlike many European countries, the country did not experience a pronounced postwar baby boom (36, 37). After nearly five decades under Soviet rule, Estonia underwent rapid institutional, economic, and educational restructuring following independence in 1991, accompanied by a sharp fertility decline and substantial postponement of childbearing (38). Such transformations have been proposed as settings in which genetic associations with socioeconomic traits may vary (39, 40). Importantly, Estonia exhibits pronounced sex differences in education–fertility associations: higher educational attainment has been associated with lower fertility among women, whereas among men education is positively associated with fertility, particularly in later birth cohorts (22, 41). These features make Estonia a well-suited setting for examining sex-specific associations between educational attainment–related polygenic scores and completed fertility.

Against this backdrop, we investigate sex-specific associations between *PGS_EA_* and completed fertility in Estonia, using data from the Estonian Biobank. We extend prior work in several ways. First, we examine a population that experienced rapid demographic and institutional change, contributing evidence beyond the Western European and US contexts that dominate existing research. Second, we utilize *PGS_Cog_* and *PGS_NonCog_* to assess whether distinct genetic EA-associated components differentially relate to fertility. Third, we evaluate the mediating and moderating roles of educational attainment and age at first pregnancy (AFP), assuming that timing of first birth is a central mechanism linking education to completed fertility. Finally, we implement within-family analyses to assess to which extent observed associations are driven by factors other than direct genetic effects.

By integrating molecular genetic data with demographic theory, our study contributes to a growing literature at the intersection of social science genomics and population research. Understanding how education-related genetic variation relates to fertility – and how these associations differ by sex and social context – clarifies the micro-level mechanisms shaping macro-level demographic change. In a period marked by sustained below-replacement fertility across much of Europe, identifying the pathways through which educational and genetic factors intersect with reproductive behavior is essential for interpreting long-run population dynamics and the evolutionary implications of contemporary demographic transitions.

## Results

We first examined population-level associations between the polygenic score for educational attainment (*PGS_EA_*) and completed fertility in the Estonian Biobank. We used a quasi-Poisson generalized linear regression model to estimate the ratio of the expected number of children, hereafter referred to as the expected fertility ratio (EFR). The EFR represents the relative multiplicative change in the expected number of children per one-unit increase in a predictor (one standard deviation for *PGS_EA_*). An EFR lower than one indicates a negative association with completed fertility, whereas an EFR greater than one indicates a positive association. Analyses were conducted separately for women (*n* = 38, 517) and men (*n* = 8, 371) who were past typical childbearing ages, as well as in a merged sample (*n* = 44, 464) (Table S2). We then assessed whether these associations differed for polygenic scores capturing the cognitive (*PGS_Cog_*) and non-cognitive (*PGS_NonCog_*) components of educational attainment, evaluated mediation by educational attainment (EA) and age at first pregnancy (AFP), and tested interactions with these factors. Finally, we examined whether associations between the polygenic scores and completed fertility were consistent with direct genetic effects.

### Associations between educational attainment polygenic scores and fertility

First, we visually assessed patterns in *PGS_EA_* across levels of completed fertility in females and males (Figure 1a and b, respectively). Among women, mean *PGS_EA_* decreased in a largely monotonic manner with increasing number of children. In contrast, mean *PGS_EA_* showed little variation across fertility groups among men.

Second, using the overall sample including both male and female individuals, we found that *PGS_EA_* is significantly negatively associated with the number of children (*EFR* = 0.974, 95% CI [0.968, 0.979], *p* = 8.5 *×* 10^−21^; Figure 1c, Table S3). However, sex-specific analyses demonstrate that this association is driven by the female subsample (*EFR* = 0.963, 95% CI [0.958, 0.969], *p* = 3.8 *×* 10^−33^), whereas the association among males is positive but not statistically significant (*EFR* = 1.007, 95% CI [0.995, 1.020], *p* = 0.25).

To assess the stability of these associations across participation waves and social environment (see Methods), we conducted additional genotype-by-environment interaction analyses. First, we found significant interaction between *PGS_EA_* and wave of participation in females (*p* = 3.4 *×* 10*^−^*^4^) that converts into a smaller negative effect of *PGS_EA_* in wave 2 (Table S4).

Next, we tested for an interaction between *PGS_EA_* and historical era (Soviet vs. post-Soviet). To operationalize historical context, individuals were classified based on their age in 1991, when Estonia regained independence from the Soviet Union. We applied two age cutoffs for women (18 or 25 years) and a single cutoff for men (25 years), reflecting the more restricted birth-year range among men. Evidence for interaction was limited. A significant interaction was observed only among women using the 25-year cutoff. However, this result was borderline after correction for multiple testing (*p* = 0.05). Consistent with this pattern, the estimated effect size of *PGS_EA_* showed some variation across birth cohorts, with an apparent increase in later cohorts, particularly among women (Figure S3a).

GWAS-by-subtraction allows the separation of genetic associations related to the cognitive and non-cognitive components of educational attainment (25). We calculated polygenic scores for the cognitive and non-cognitive components of EA (*PGS_Cog_* and *PGS_NonCog_*, respectively) using the corresponding GWAS summary statistics. Consistent with the results for *PGS_EA_*, both *PGS_Cog_* and *PGS_NonCog_* were negatively associated with completed fertility among women and showed no significant associations among men (Figure 1d, Table S5). Despite a relatively small difference in their correlations with *PGS_EA_* (0.44 vs. 0.49, females; Table S2), the association with female completed fertility was significantly stronger for *PGS_NonCog_* than for *PGS_Cog_* (*p*-value for difference = 1.1 *×* 10*^−^*^3^).

### Mediation and moderation effects of educational attainment

EA is a potential mediator of the relationship between *PGS_EA_* and completed fertility. To test this, we included EA in the regression model alongside *PGS_EA_*. We considered two alternative EA measures: a finely defined measure consisting of 11 education levels, and alternatively a simplified phenotype with three ordered education categories treated as a continuous variable (basic = 0, intermediate = 1, and advanced = 2) (Table S1). In both specifications, the estimated effect size of *PGS_EA_* in women was 30% smaller than in the model without EA (Figure 2a, Table S6). Given the similarity of results, we used the three-category EA measure in subsequent analyses for ease of interpretation.

**Figure 1:**
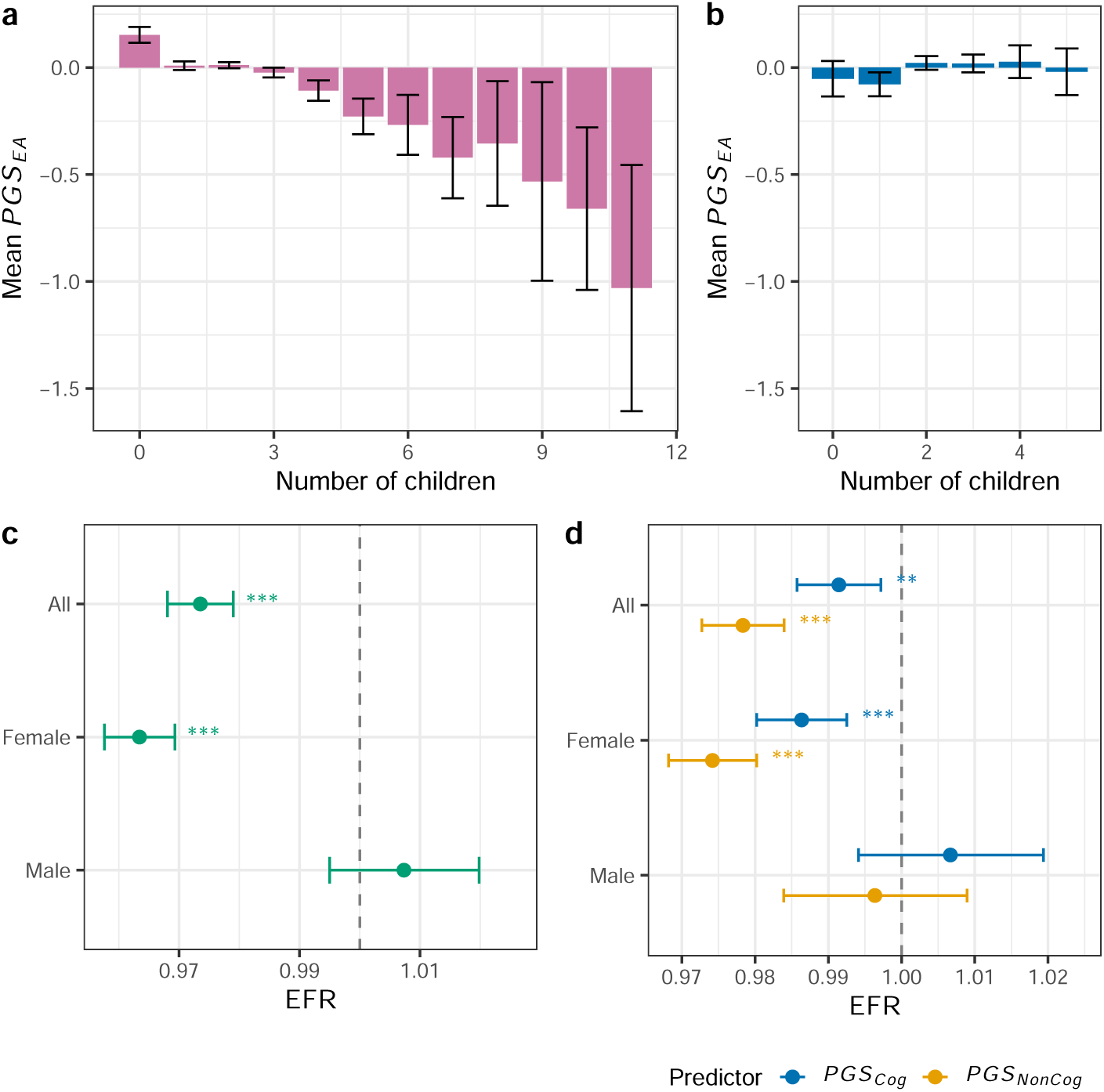
Population-level associations between educational attainment polygenic scores and fertility. Average *PGS_EA_* values and their 95% confidence intervals among females (a) and males (b) in groups by number of children. For males, the number of children “5” corresponds to “5 or more”. Expected fertility ratios (EFR) for the association between (c) *PGS_EA_* or (d) *PGS_Cog_*/*PGS_NonCog_* and number of children in unrelated individuals, shown for the full sample and stratified by sex. All models adjust for year-of-birth categories, the first 10 genetic principal components, county of birth, and participation wave. EFRs for *PGS_Cog_* and *PGS_NonCog_* were estimated jointly in the same model. The error bars denote 95% confidence intervals. The statistical significance level is denoted by asterisks (* *P <* 0.05, ** *P <* 0.01, *** *P <* 0.001).

The three-category EA measure was significantly associated with completed fertility in both females and males. The effect was negative among females but positive among males. Including EA in the model also attenuated the associations of *PGS_Cog_* and *PGS_NonCog_* with completed fertility among females (Figure 2b, Table S7), rendering the association for *PGS_Cog_* non-significant. Consistent with this pattern, in linear models with EA as the outcome, *PGS_Cog_* showed a non-significantly but stronger association with EA than *PGS_NonCog_* (*PGS_Cog_*: *β* = 0.111, SE = 0.003; *PGS_NonCog_*: *β* = 0.103, SE=0.003; *p*-value for difference = 0.034).

Previous studies have shown that socioeconomic status (SES) in general, and EA in particular, can moderate the association between *PGS_EA_* and fertility (14, 15). We tested this by including the interaction term *EA × PGS_EA_* in the regression model (Table S8). Its effect was significantly positive in women (*p* = 7.5 *×* 10^−19^), counteracting the negative main effects of EA and *PGS_EA_*.

To further explore potential non-linearities in the interaction between EA and *PGS_EA_*, we stratified the female and male cohorts by EA (Figure 2c, Table S9) and *PGS_EA_* (Figure 2d, Table S10) and estimated the effects of *PGS_EA_* and EA within these strata, respectively. Among women, the effect of *PGS_EA_* was strongest among individuals with basic education (*EFR* = 0.925, 95% CI [0.891, 0.959], *p*= 3.2*×*10*^−^*^5^), followed by those with intermediate education (*EFR* = 0.961, 95% CI [0.952, 0.970], *p* = 5.7 *×* 10^−17^). The effect was still significant but substantially weaker in women with advanced education (*EFR* = 0.991, 95% CI [0.983, 1], *p* = 0.046).

The effect of EA was substantially stronger in female strata with low *PGS_EA_* values than in strata with higher *PGS_EA_* values. The estimates for males were relatively stable, consistent with no interaction found in the linear model.

Variation in the estimated effect size of *PGS_EA_* across EA cohorts may partly explain the observed interactions with participation wave and historical era demonstrated above. Wave 2 participants represent a population cohort with a higher average level of educational attainment than wave 1 participants (42). Average educational attainment is also higher in more recent birth cohorts. Consistent with this interpretation, including EA and the interaction term *PGS_EA_×EA* in the corresponding models rendered the initial interactions with participation wave and historical era non-significant (Table S4). In addition, in stratified analysis by education level, differences in the estimated *PGS_EA_* effect sizes across birth-year cohorts were substantially reduced (Figure S3bc), confirming that the temporal trend was cased by increase in average education level.

### Mediation and moderation effects of age at first pregnancy

To explore the role of age at first pregnancy in the relationship between *PGS_EA_* and completed fertility, we repeated the mediation and moderation analyses using AFP, restricting the sample to women who had ever been pregnant.

AFP is a strong predictor of completed fertility, with the effect of a one-year increase in AFP comparable to that of a one–standard deviation increase in *PGS_EA_* (Figure 3a, Table S11). Including AFP as a covariate substantially attenuated the association between *PGS_EA_* and completed fertility (*EFR* = 0.996, 95% CI [0.991, 1.002], *p* = 0.214), consistent with reproductive timing mediating this relationship. Interestingly, while AFP had a similar attenuating effect on *PGS_NonCog_*, it changed the estimated effect of *PGS_Cog_* to positive (*EFR* = 1.008, 95% CI [1.002, 1.014], *p* = 6.2 *×* 10*^−^*^3^).

**Figure 2:**
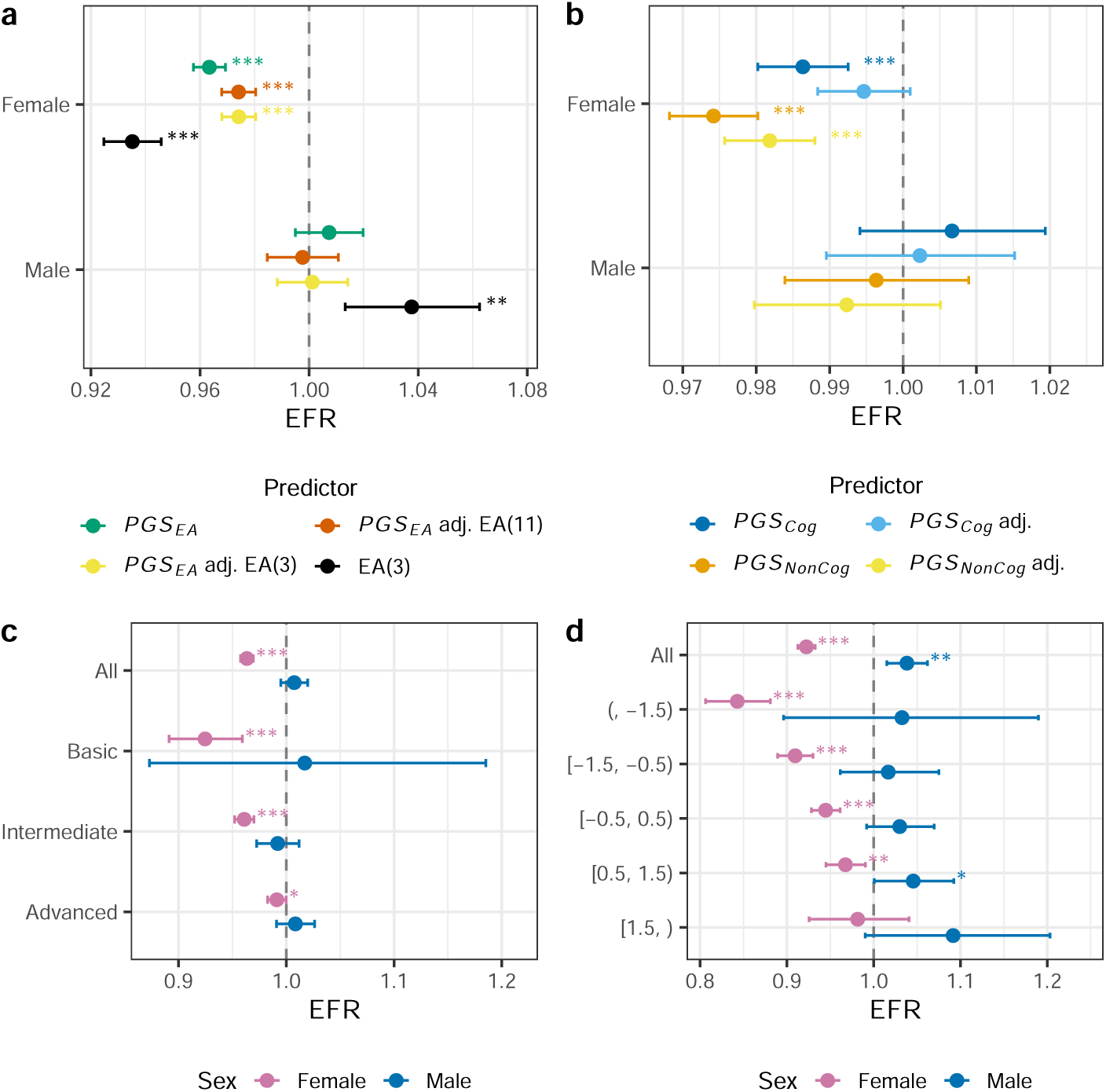
Educational attainment as mediator and moderator of genetic effects on fertility. Expected fertility ratios (EFRs) for the association between polygenic scores or educational attainment (EA) and number of children in unrelated individuals. (a) Associations between *PGS_EA_* and fertility before and after adjustment for EA, modeled either as a categorical variable with 11 levels or as an ordinal variable with 3 levels, shown separately for women and men. (b) Associations of *PGS_Cog_* and *PGS_NonCog_* with fertility before and after adjustment for ordinal EA, alongside the association between EA and fertility. (c) Associations between *PGS_EA_* and fertility stratified by EA category (basic, intermediate, advanced), shown separately for women and men. (d) Associations between EA and fertility stratified by *PGS_EA_* categories, shown separately for women and men. Models adjust for year-of-birth categories, county of birth, wave of participation, and the first 10 genetic principal components. The error bars denote 95% confidence intervals. The statistical significance level is denoted by asterisks (* *P <* 0.05, ** *P <* 0.01, *** *P <* 0.001).

In a joint regression model including EA and AFP alongside *PGS_EA_*, the EFR for EA is substantially closer to one than in the model including EA alone (0.980, 95% CI [0.970, 0.990] vs. 0.935, 95% CI [0.925, 0.946], respectively), whereas the EFR for AFP is largely unchanged compared with the model including AFP alone (Table S12). This pattern suggests that the association between EA and fertility mainly reflects the correlation between EA and AFP. Notably, in the joint model, a one-year delay in AFP is associated with a larger fertility reduction than the contrast between adjacent education categories (0.970, 95% CI [0.969, 0.972] vs. 0.980, 95% CI [0.970, 0.990], respectively).

The interaction term *AFP × PGS_EA_* was significantly positive (*p* = 2.3 *×* 10*^−^*^9^; Table S8). In analyses stratified by AFP, the effect of *PGS_EA_* was negative among individuals with AFP *<* 20, approximately zero among those with 20 *≤* AFP *<* 25, and positive among those with AFP *≥* 25 (Figure 3b, Table S13). Further adjustment for AFP within strata did not substantially alter the estimates, indicating that the moderating effect of AFP operates primarily across AFP strata rather than through residual within-stratum variation. The effect of AFP was negative in all strata defined by *PGS_EA_* and was generally stronger in groups with lower *PGS_EA_* (Figure 3c, Table S14).

**Figure 3:**
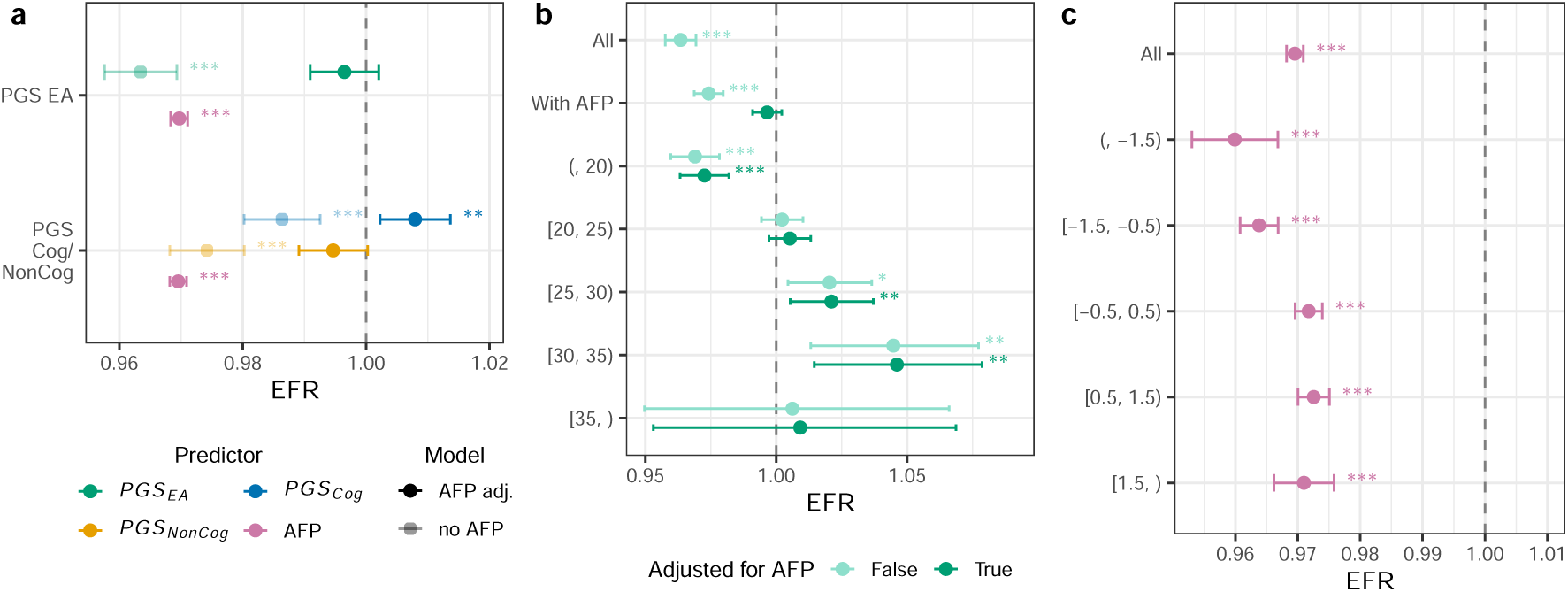
Age at first pregnancy as mediator and moderator of genetic effects on fertility in women. Expected fertility ratios (EFRs) for the association between polygenic scores or age at first pregnancy (AFP) and number of children in unrelated women ever been pregnant. (a) Associations of *PGS_EA_*, *PGS_Cog_*, and *PGS_NonCog_* with fertility before and after adjustment for AFP; increased transparency indicates models without AFP adjustment. (b) Associations between *PGS_EA_* and fertility stratified by AFP categories among women who have ever experienced pregnancy. “All” states for the complete unrelated female sample regardless of the AFP phenotype availability, while “With AFP” indicates the subsample of unrelated women for which the AFP information is available. Models were estimated with and without additional adjustment for AFP within each stratified group. (c) Associations between AFP and fertility stratified by *PGS_EA_* categories among women. Models adjust for year-of-birth categories, county of birth, wave of participation, and the first 10 genetic principal components. The error bars denote 95% confidence intervals. The statistical significance level is denoted by asterisks (* *P <* 0.05, ** *P <* 0.01, *** *P <* 0.001).

### Within-family genetic effects on fertility

The effects of *PGS_EA_* represent the sum of dynastic effects, effects of population structure and assortative mating, as well as direct genetic effects (26, 43). To examine, whether the effects of *PGS_EA_* on completed fertility include a direct genetic component we utilized family based designs.

First, we used paternal and maternal polygenic scores for EA derived from imputed and genotyped parental genotypes. Including parental polygenic scores as covariates in the regression models allows correction for indirect genetic effects, assortative mating and residual population structure, thereby enabling estimation of the direct (within-family) genetic effects of the PGS. In models including parental *PGS_EA_*, we used an alternative individual *PGS_EA_* (proband *PGS_EA_*), constructed using the same set of genetic variants as the parental polygenic scores to ensure unbiased adjustment for direct genetic effects. Consistent with previous analyses, we used a quasi-Poisson generalized linear regression model. One individual per sibship was randomly selected for inclusion in the sample.

The population-level effects of the proband *PGS_EA_* were significantly negative among females and significantly positive among males (Figure 4a; Table S15). Within-family *PGS_EA_* effect estimates were larger in absolute magnitude than the corresponding population-level estimates and remained significantly different from zero in both sexes. These results are consistent with the population-level associations between *PGS_EA_* and completed fertility being driven predominantly by direct genetic effects. To assess the robustness of these results, we alternatively used the linear mixed model implemented in snipar (44). This model treats sibship as a random effect, allowing the inclusion of siblings, and accounts for assortative mating, thereby reduc-ing bias in parental PGS effect estimates. Despite differences in model specification, results from the quasi-Poisson and snipar models were largely consistent (Figure S4). Parental effect estimates were not significantly different from zero.

Second, we applied a sibling design to access within-family effects of *PGS_EA_*. Owing to the small male sample size, this analysis was conducted only in women. We estimated fixed-effects models including the mean sibship *PGS_EA_* (corresponds to between-family effect) and individual deviations from this mean (corresponds to within-family effect). All three estimates (population, within- and between-family) were negative and similar in magnitude, with the within-family point estimate slightly more negative than the others (Figure 4b, Table S16).

For *PGS_Cog_* and *PGS_NonCog_*, within-family estimates could not be distinguished from either the population estimates or zero because of limited statistical power. Nevertheless, the within-family point estimates were again more extreme than the corresponding population and between-family estimates. Taken together, these findings further support the interpretation that the observed population associations are largely attributable to direct genetic effects.

Given the potential mediating roles of EA and AFP in the association between *PGS_EA_* and completed fertility, we repeated the family-based analyses for these phenotypes using linear mixed models implemented in snipar (44).

For EA, the effect sizes of the proband *PGS_EA_* adjusted for parental *PGS_EA_* were attenuated relative to the corresponding population-level effects in both sexes. The difference between the population and within-family effects was statistically significant among females (*p* = 3.7 *×* 10*^−^*^7^; Figure S5a, Table S18). The effects of both maternal and paternal *PGS_EA_* were significant among females, whereas only the paternal effect was significant among males. One should note, however, that parental PGS effects are biased towards their mean because parental genotype imputation using a sibling genotype cannot distinguish the parent of origin. Consistent with this pattern, within-family effects in the analysis of siblings were significantly smaller than between-family effects among women (*p* = 3.2 *×* 10*^−^*^7^; Figure S5c, Table S19).

**Figure 4:**
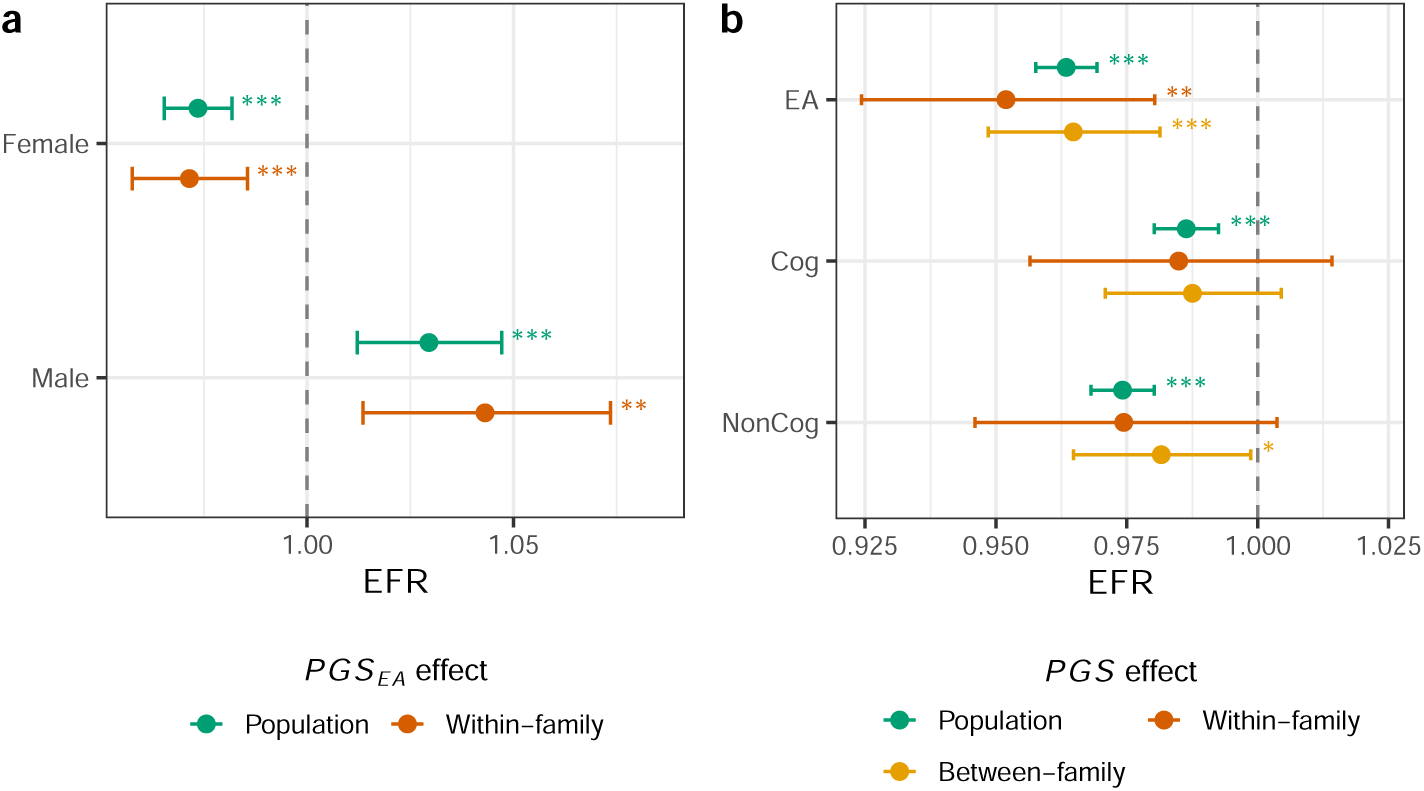
Within-family genetic effects on fertility. Expected fertility ratios (EFRs) for the association between polygenic scores and number of children. (a) Population-level and within-family associations between *PGS_EA_* and completed fertility. “Within-family” effect corresponds to the regression coefficient of individual *PGS_EA_* adjusted for parental *PGS_EA_*. The individual (proband) *PGS_EA_* was constructed using the same set of SNPs as the parental *PGS_EA_*. (b) Decomposition of *PGS* effects on fertility in women into population, within-sibling, and between-sibling components, shown for *PGS_EA_*, *PGS_Cog_*, and *PGS_NonCog_*. Population models were estimated in unrelated individuals. All models adjust for year-of-birth categories, county of birth, wave of participation, and the first 10 genetic principal components. The error bars denote 95% confidence intervals. The statistical significance level is denoted by asterisks (* *P <* 0.05, ** *P <* 0.01, *** *P <* 0.001).

A similar pattern was observed for AFP. The proband *PGS_EA_* effect on AFP, adjusted for parental *PGS_EA_*, was significantly smaller than the population-level effect (*p* = 4.0 *×* 10*^−^*^3^; Figure S5b, Table S20), and the effects of both maternal and paternal *PGS_EA_* were significant. In the sibling analysis, within-family effects were again significantly smaller than between-family effects (*p* = 8.6 *×* 10*^−^*^3^; Figure S5d, Table S21).

## Discussion

This study examines sexand environment-specific associations between polygenic scores for educational attainment and completed fertility in Estonia, broadening the geographic, social, and genetic scope of genetically informed fertility research. By utilizing polygenic scores for cognitive and non-cognitive components of educational attainment, we contribute to disentangling the mechanisms underlying these associations. Leveraging family-based designs, we further distinguish pathways consistent with direct genetic effects from those driven by correlated environmental factors.

In line with the few existing studies examining associations between *PGS_EA_* and fertility in the US (16), Iceland (13), and the UK (45), we observed a negative association among women. In contrast to these studies, the association among men in Estonia was non-significant or even positive, depending on the PGS considered.

The directions of these sex-specific associations mirror the patterns between educational attainment and fertility in Estonia reported in demographic studies (22, 41) and observed in our own analysis. Prior research across multiple societies has also documented opposing associations between socioeconomic status and fertility in men and women (3, 46, 47). However, this pronounced sex difference stands in contrast to expectations based on the near-perfect genetic correlation of fertility between sexes (34). The observed reduction in female fertility may instead reflect EA-associated variants operating under specific environmental conditions, with effects that are too context-dependent or heterogeneous to be detected in GWAS meta-analysis of fertility, or arise from non-causal genotype–environment correlations.

Our results indicate that educational attainment plays only a modest role in mediating the association between *PGS_EA_* and fertility. First, the EA phenotype explains only a fraction of the association between *PGS_EA_* and fertility among women. Second, we found a stronger negative association between female fertility and *PGS_NonCog_* compared to *PGS_Cog_*, even though *PGS_Cog_* is a marginally stronger predictor of educational attainment. This finding contrasts with the largely indistinguishable genetic correlations between fertility and the cognitive and non-cognitive components of educational attainment (25). Third, adjustment for EA attenuated the association of *PGS_Cog_* with fertility more strongly than that of *PGS_NonCog_*. This finding underscores the importance of non-cognitive characteristics in shaping fertility beyond education level. Discrepancies with other studies (48, 49) may reflect their insufficient statistical power to detect *PGS_EA_* effects beyond those mediated by intelligence and education. Fourth, the moderate association between EA and fertility was substantially attenuated in the joint model with AFP, whereas the effect of AFP was largely unchanged. Fifth, we found no evidence for indirect genetic effects of *PGS_EA_* on fertility. This result is consistent with comparable population-level and within-family genetic correlations between EA and fertility (35). At the same time, we observe substantial indirect effects of *PGS_EA_* on EA.

A plausible explanation reconciling the negative association between educational attainment and fertility with these observations is that both outcomes are influenced by intermediate traits, such as personality, partnership formation, or life values, that have a genetic component captured by *PGS_EA_* (2). This interpretation is supported by prior demographic studies showing that the direct effect of EA on fertility among women may be close to zero or even positive, in contrast to the negative association observed at the population level (50, 51).

Interactions between *PGS_EA_* and educational attainment in the context of fertility have been reported previously. For example, the association between *PGS_EA_* and fertility was weaker among individuals, especially men, with higher levels of education (13, 14). At the phenotypic level, the fertility-reducing effect of higher education was strongest among women from disadvantaged social backgrounds and with lower early achievement (52).

One explanation is that attaining higher education from a disadvantaged starting point requires greater time and effort, thereby intensifying the trade-off between education or career and childbearing. Such disadvantages may reflect the environmental as well as genetic background (14, 15). Alternatively, the interaction between the EA level and *PGS_EA_* may reflect a mismatch between genetic predisposition (direct or indirect) and realized educational attainment. High *PGS_EA_* combined with low attainment may indicate unmeasured psychological, environmental, or genetic constraints affecting both education and fertility, whereas low attainment driven primarily by low *PGS_EA_* or correlated factors may have weaker fertility consequences. In addition, because highly educated women tend to have fewer children and lower variance in fertility, there may be limited scope for *PGS_EA_* effects to manifest.

The interaction between *PGS_EA_* and educational attainment also accounts for the observed interactions between *PGS_EA_* and historical era or participation wave. Despite substantial societal changes during the study period, the association between *PGS_EA_* and fertility appears largely stable once differences in educational attainment are taken into account.

In our study, age at first pregnancy accounted for much of the association between *PGS_EA_* and fertility, in agreement with previous observations (13). However, AFP explained the *PGS_EA_* effect only on average. In analyses stratified by AFP, the association between *PGS_EA_* and fertility varied across groups and changed direction, from negative in the lowest-AFP group to positive in high-AFP groups. This observation is consistent with prior findings (13, 14). Meanwhile, later reproductive onset was associated with lower fertility across all *PGS_EA_* strata, but this association was strongest among women with lower *PGS_EA_*.

Several complementary mechanisms may underlie these patterns. First, they may reflect within-population demographic trajectories that parallel those observed across countries, whereby fertility transitions occur first in higher-SES strata, just as they occur first in more economically developed societies (53). Early in the transition, late births decline as individuals limit family size. Subsequently, late births increase as childbearing is postponed (10, 54). If this transition diffuses gradually across social strata, higher-SES groups may postpone reproduction while maintaining longer reproductive windows, whereas lower-SES groups may curtail reproduction earlier. Within this framework, *PGS_EA_* captures residual SES-related heterogeneity in AFPstratified analyses: among women who initiate childbearing later, those with higher *PGS_EA_* ultimately achieve higher completed fertility. Second, *PGS_EA_*, and particularly *PGS_NonCog_*, may shape how reproductive timing interacts with career orientation. Among women who begin childbearing early, stronger career orientation may intensify the trade-off between continued childbearing and educational or occupational investment, resulting in lower completed fertility. In contrast, among women who postpone childbearing, higher *PGS_EA_* may be associated with greater economic and career stability, allowing higher fertility once reproduction begins. Third, later reproductive onset among individuals with lower *PGS_EA_* may reflect persistent constraints rather than strategic postponement. Health limitations, socioeconomic disadvantage, or other adverse conditions may delay entry into parenthood while simultaneously limiting fertility at later ages. In contrast, delayed childbearing among individuals with higher *PGS_EA_* is more likely to reflect postponement compatible with extended reproductive capacity. Together, these mechanisms suggest that *PGS_EA_* differentiates between delayed reproduction arising from constraint versus postponement consistent with longer reproductive windows and higher completed fertility.

By applying family-based models, we were able to show that most of the association between *PGS_EA_* and fertility in women can be explained through direct genetic effects. Despite being direct, these effects may operate through intermediate traits, such as personality, active or reactive genotype-environment correlations (55). Furthermore, direct genetic effects may be modified by genotype-by-environment interactions, implying that changes in the social environment could mitigate their impact. These findings motivate further research on whether social contexts differentially shape reproductive opportunities for individuals with different genetic predispositions.

Our study has several limitations. Although participants’ birth years span more than five decades, most were born between 1945 and 1975, limiting long-term extrapolation. Age at first pregnancy was used as a proxy for age at first birth due to data availability; while closely related, these measures are not identical. The sample size for men was substantially smaller than for women because of a stricter age threshold, lower participation rate, and more limited questionnaire coverage, reducing statistical power. Participation and mortality biases, particularly in earlier birth cohorts, lead to overrepresentation of individuals with higher education and better health, which is expected to attenuate estimated effects among women. Although family-based designs reduce confounding by shared environment and population stratification, they do not eliminate all sources of bias (56, 57), and causal interpretations should remain cautious. Finally, the observed patterns may reflect context-specific features of Estonia’s demographic and institutional history, underscoring the need for broader geographic coverage to assess generalizability.

Future studies will require larger sample sizes to disentangle higher-order interactions and to examine how cognitive and non-cognitive genetic components of educational attainment interact with environmental contexts. Family-based cohort designs will remain essential for strengthening inference about direct genetic effects. Broader measurement of socioeconomic, psychological, and life-course traits will be critical for developing a more comprehensive understanding of the mechanisms shaping fertility in contemporary populations.

By combining population-level, stratified, and family-based analyses in a rapidly transforming Estonian society, this study advances understanding of how educationrelated genetic variation relates to fertility in a sex-specific and context-dependent manner. Our results suggest that EA-related polygenic scores capture broader life-course propensities beyond education itself, and that their fertility consequences cannot be reduced to linear mediation through schooling or reproductive timing. Future work in larger and more diverse populations will be essential for disentangling these pathways and clarifying how genetic influences on fertility interact with social change.

## Materials and Methods

### Participants

Study participants were a subsample of the Estonian biobank (EstBB), a population-based cohort of approximately 210,000 adult individuals (*∼*138,000 women and *∼*73,000 men) residing in Estonia (42, 58). Participants were born between 1905 and 2003 (mean, 1972; SD, 16.5). They were recruited between 2001 and 2021 in two waves (wave 1: from 2001 to 2012; wave 2: from 2017 to 2021). During the first wave, partic ipants were mostly recruited by general practitioners, while a broad media campaign took place during wave 2. The data collected includes genotype, demographic, health, and lifestyle information and blood samples.

### Ethics statement

The activities of the EstBB are regulated by the Human Gene Research Act, which was adopted in 2000 specifically for the operations of the EstBB. Individual-level data analysis in the EstBB was carried out under ethical approval “1.1-12/2566” from the Estonian Committee on Bioethics and Human Research (Estonian Ministry of Social Affairs), using data according to release application “6-7/GI/10508” from the Estonian Biobank.

### Genotyping and quality control

Genotyping was performed using the Infinium Global Screening Arrays (GSA) with approximately 550,000 variants overlapping across the different versions of the plat form. Individuals with a call rate *<* 95% or a mismatch between chromosomal and self-reported sex were excluded from the analysis. Only strand-unambiguous SNPs were kept for the imputation. Next, the genotypes were imputed with Beagle 5.4 (59) using the EstBB Estonian-specific reference panel.

### Ancestry and PCA

Genetic ancestry grouping was estimated with bigsnpr (60) on genotypes imputed using 1000 Genomes Project phase 3 samples (61) for better representation of global haplotype diversity. Further analyses were restricted to individuals of European ancestry (“East European”, “North-West European”, or “Finnish”).

Principal component analysis (PCA) was performed using flashPCA (62) to capture population structure. PCA was first conducted in a subset of individuals after excluding related individuals up to third-degree relatedness. The analysis was based on approximately 65,000 biallelic SNPs pruned for linkage disequilibrium using the 1000 Genomes Project Phase 3 reference panel (61). Individuals with inferred relatedness were subsequently projected onto the resulting principal component axes. Samples with values exceeding 7 standard deviations from the mean on any of the first 10 principal components were excluded. PCA was then repeated in the cleaned unrelated subset, and related individuals were again projected onto the final principal component coordinates.

### Phenotype construction

#### Fertility

The primary outcome was completed fertility, defined as the total number of children ever born. Fertility information was derived from two independent sources: (i) detailed reproductive histories from the women’s health questionnaire and (ii) a later personality questionnaire reporting the total number of children (63).

For women, completed fertility was derived using both sources. Based on the women’s health questionnaire data, it was defined as the minimum of the last reported number of pregnancies and the last reported number of live births. In most cases, the number of pregnancies was equal to or greater than the number of live births. The personality questionnaire allowed participants to report the number of children as “1,” “2,” “3,” “4,” or “5 or more.” For women, responses of “5 or more” were treated as missing because an exact count could not be recovered reliably. When fertility information was available from both questionnaires, observations were excluded if the absolute difference between sources exceeded one child. The final fertility measure was defined as the maximum of the harmonized reproductive-history count and the personality questionnaire count, allowing for child mortality or births occurring after the reproductive-history survey.

For men, fertility information was obtained from the personality questionnaire. To maximize sample size, responses of “5 or more” children were interpreted as “5.” Analyses were restricted to individuals who were past typical childbearing ages: women older than 45 years and men older than 50 years at the time of assessment.

#### Age at first pregnancy

Age at first pregnancy (AFP) was derived from women’s reproductive histories. When multiple values were reported, the mean value was used. For the interaction analysis, AFP was centered by subtracting the sample mean (22.6 years). We use AFP as a proxy for age at first life birth, as the information on the latter is not available for the EstBB participants.

#### Educational attainment

Educational attainment (EA) was obtained from registry-linked education codes as well as from various questionnaires. The highest reported level of education was used. For specific analyses, EA was modeled either as (i) an 11-level categorical variable or (ii) a continuous variable with three possible values (basic = 0; intermediate = 1; advanced = 2) (Table S1).

#### Covariates

In all regression models, ten genetic principal components were included as covariates to account for population structure. Year of birth was included as a categorical covariate using dummy variables, allowing us to adjust for nonlinear changes in the mean outcome over time without specifying a parametric functional form.

Previous studies have shown that passive genotype–environment correlations can inflate genotype–phenotype associations (45), and that such correlations are present in contemporary Estonia (64). To further reduce potential effects of population structure, we included county of birth as an additional covariate in the regression models, alongside ten genetic principal components.

The EstBB recruitment process consisted of two major waves, with most participants recruited in 2004–2010 and 2018-2022. These waves differed in recruitment strategy and therefore represent differentially biased population samples (42). To mitigate potential inter-wave effects on our estimates, we also included wave of participation (wave 1: 2001-2016; wave 2: 2017-2022) as a covariate in the model.

After adjusting for county of birth and wave of participation, estimates changed modestly (Figure S2, Table S3).

### Polygenic scores

The primary polygenic score for educational attainment (*PGS_EA_*), proband *PGS_EA_* and parental *PGS_EA_* were obtained from the PGI Repository (65). These *PGS* were constructed using summary statistics from the most recent genome-wide association study (EA4) (12), with the Estonian Biobank cohort excluded from the meta-analysis. Polygenic score weights were reestimated using SBayesR (66).

Parental *PGS_EA_* were derived from parental genotypes imputed using Mendelian imputation (44). Because quality control criteria for imputed parental variants were more stringent than those applied for genotyped individuals, a smaller set of variants was available for parental *PGS_EA_* construction. To ensure consistency, an alternative proband *PGS_EA_* based on the same variant set was used in analyses that included parental *PGS_EA_*.

In addition, separate polygenic scores capturing the cognitive (*PGS_Cog_*) and noncognitive (*PGS_NonCog_*) components of educational attainment were calculated using summary statistics from GWAS-by-subtraction analysis, again excluding the Estonian Biobank cohort (25).

All polygenic scores were standardized to have a mean of zero and unit variance.

### Identification of unrelated individuals and sibships

Relatedness between individuals was inferred with KING (67). Individuals with 2nddegree relatedness or closer were excluded from the main analyses. Sex-specific unrelated samples were created for men and women, as well as a combined unrelated sample. Overall, 46,564 unrelated individuals were left in the combined sample, 40,738 females and 8,371 males in the respective sex-specific samples (Figure S1). The combined sample size is smaller than the sum of the female and male sample sizes because individuals related across sexes could be included in the sex-specific analyses but were excluded from the combined analysis.

Sibling-based female sample was used for within-family analyses, comparing female siblings within the same family, thus controlling for shared family background. Siblings were defined by KING criteria of full siblings and an additional threshold of 0.177 *< Kinship <* 0.354. Only full sibships were considered (in which all members were classified as siblings).

### Statistical analysis

#### Population-level analysis

Population-level associations between fertility and genetic predictors were estimated using quasi-Poisson generalized linear regression models with a log link. The outcome variable was the number of children per individual. In the main analysis, the conditional mean of completed fertility was modeled as a function of the educational attainment polygenic score, participation wave, county of birth fixed effects, birthyear categorical fixed effects, and the first ten genetic principal components (eq. (1)). The quasi-Poisson specification allows the variance of the outcome to differ from the mean by a constant dispersion factor, relaxing the Poisson assumption while preserving the same conditional mean structure. This approach yields consistent coefficient estimates while adjusting standard errors for the observed underdispersion in the number of children. Exponentiated coefficients (expected fertility ratios, EFRs) are interpreted as multiplicative changes in the expected number of children associated with a one-unit increase in the corresponding covariate. Analyses were conducted in the full unrelated sample and stratified by sex.

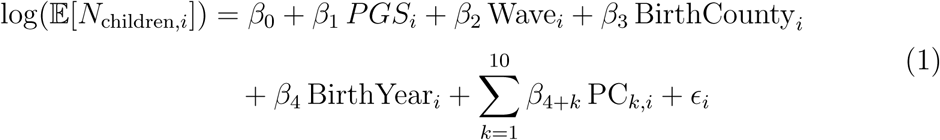

Here *N*_children_*_,i_* denotes the number of children of individual *i*. *PGS_i_* denotes the educational attainment polygenic score of individual *i*. Wave*_i_* denotes the participation wave, for which the values 0 and 1 correspond to wave 1 and wave 2, respectively. BirthCounty*_i_* denotes fixed effects county of birth, and BirthYear*_i_* denotes fixed ef-fects categorical year of birth. PC*_k,i_*, where k=[1,10], denote first ten genetic principal components. The residual is represented as *ɛ_i_*.

In mediation analyses, regression models additionally included educational attainment or age at first pregnancy as covariates.

To examine linear statistical interactions between *PGS_EA_* and EA, AFP, wave of participation, or historical era, we included the corresponding main effects and interaction terms in the regression models. For analyses involving participation wave or historical era, we also estimated models that additionally adjusted for EA and included the interaction term *PGS_EA_ × EA*.

To assess potential non-linear interactions between *PGS_EA_* and EA or AFP, we conducted stratified analyses across categories of EA, AFP, or *PGS_EA_*. In analyses stratified by AFP, models additionally adjusted for AFP within strata to account for residual within-stratum variation.

*PGS_Cog_* and *PGS_NonCog_* were included jointly when estimating their independent associations.

#### Within-family effect analysis using parental PGS

Two models were used in the within-family analysis of parental *PGS_EA_*. The first model, applied to completed fertility, was a fixed-effects quasi-Poisson generalized linear regression model in which one individual per sibship was randomly selected for inclusion. Maternal and paternal *PGS_EA_* were included as covariates. This model does not correct for assortative mating and therefore does not provide unbiased estimates of parental genetic effects. However, it is consistent with the model used in the fertility analyses elsewhere in this study.

The second model was implemented in snipar v0.0.18 and is recommended for use with imputed parental PGS (44). snipar fits a linear mixed model with sibship treated as a random effect and includes correction for assortative mating, thereby enabling less biased estimation of parental genetic effects. This model was used for robustness analyses of completed fertility, as well as for educational attainment (EA) and age at first pregnancy (AFP).

#### Within-family effect analysis using sibling design

Within-family models were estimated using sibling fixed effects, which decompose genetic associations into within-sibling and between-sibling components (eq. (2)). The sibship-mean *PGS* was used to estimate between-sibship effects, while individual deviations from the sibship mean were used to estimate within-sibship effects in a joint regression model. We assume no sibling indirect genetic effects. Under this assumption, the within-sibship estimates correspond to direct genetic effects, whereas the between-sibship estimates capture the sum of direct effects and non-direct genetic effects, such as dynastic effects, residual population stratification, and assortative mating.

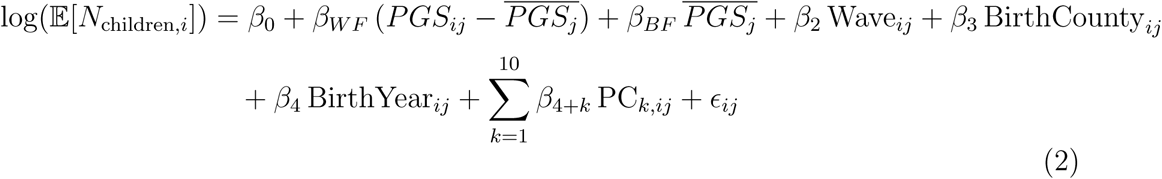

Here, *N*_children_*_,i_* denotes the number of children of individual *i* from sibship *j*. *PGS_ij_* denotes the educational attainment polygenic score of individual *i* from sibship *j*, and 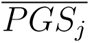 denotes the mean *PGS_EA_* of sibship *j*. All covariates correspond to the regression terms from eq. (1), with the difference that the index *ij* indicates individual *i* from sibship *j*. The coefficients *β_W F_* and *β_BF_* represent the within-family and between-family effect estimates, respectively.

#### Statistical significance

Testing whether regression coefficients differed from zero was performed using two-sided t-test. The difference between coefficients from the same model (*PGS_Cog_* and *PGS_NonCog_*) was tested with a Wald test. A threshold of *α* = 0.05 was used for statistical significance. 95% confidence intervals are reported throughout unless otherwise stated. In analyses involving multiple age cutoffs, Bonferroni correction was applied to adjust for multiple testing.

#### Software

Statistical analyses were conducted in R v4.4 (68) using custom code, including the lme4 package (69). Figures were generated using the ggplot2 package (70).

### Data availability

Access to the Estonian Biobank data (https://genomics.ut.ee/en/content/estonian-biobank) is restricted to approved researchers and can be requested.

## Code availability

Custom R code used for statistical analyses is publicly available on GitHub (https://github.com/ivkuz/PGSeaFertility).

## Acknowledgments

The authors acknowledge the participants of the Estonian Biobank. We thank Tobias Edwards for fruitful discussions, René Moñttus for his effort in designing the personality questionnaire and collecting the corresponding data, Francesca Procopio for providing us with summary statistics for cognitive and non-cognitive components of educational attainment. The research was conducted using the Estonian Center of Genomics/Roadmap II, funded by the Estonian Research Council (project number TT17). Data analysis was carried out in part in the High-Performance Computing Center of the University of Tartu. IK was supported by the Centre of Excellence for Personalised Medicine, funded by grant TK214 from the Estonian Ministry of Education and Research. AG and CR were supported by the European Research Council [GEPSI 946647]. The funders had no role in the study design, data collection and analysis, decision to publish, or preparation of the manuscript. The personality study has been funded by Estonian Research Council’s personal research funding start-up grants PSG656 and PSG759, and Estonian Research Council’s team grants PRG2190 and PRG1291.

## Author contributions

I.K. conceptualized the study, conducted the statistical analyses, interpreted the results, and wrote the initial draft. A.G. contributed to study conceptualization, interpretation of the results, and writing the initial draft. K.L., C.R., U.V., and V.P. contributed to study design and interpretation. All authors reviewed and edited the manuscript. The Estonian Biobank Research Team and U.V. contributed data and infrastructure.

