## Supplementary Figures and Tables for "Sex-specific associations between education-related genetic factors and fertility extend beyond educational attainment"

### Supplementary Materials

Ivan A. Kuznetsov, Alexandros Giannelis, Estonian Biobank Research Team, Kelli Lehto, Triin Laisk, Cornelius A. Rietveld, Uku Vainik, Vasili Pankratov

### Supplementary figures

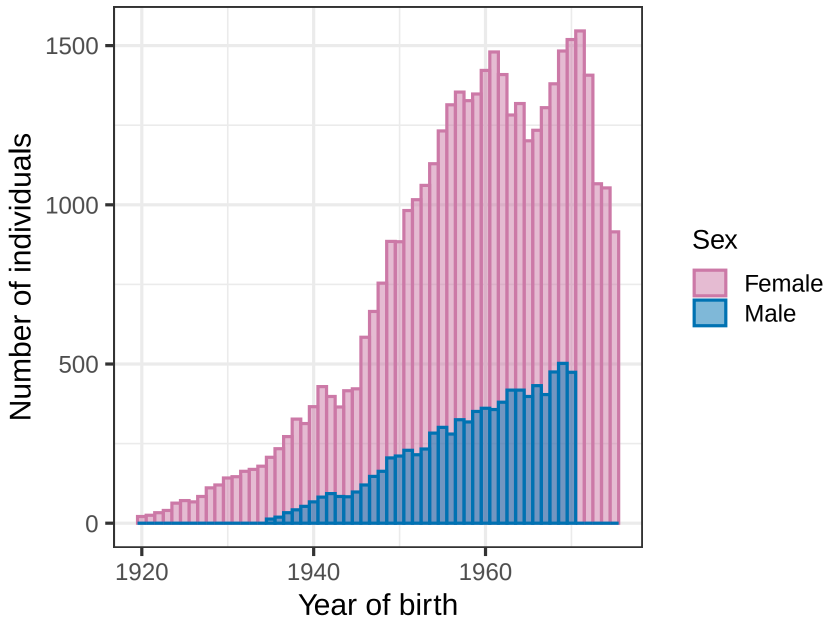


Figure S1: **Sample distribution by sex and year of birth.**

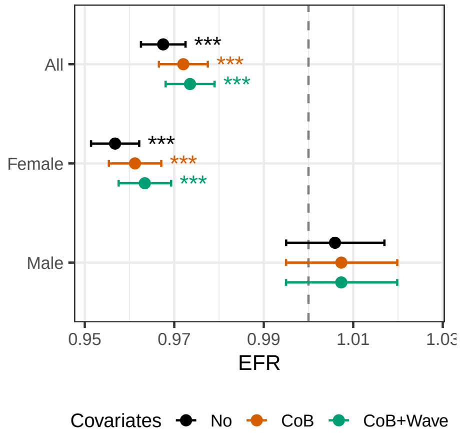


Figure S2: **Population-level associations between $PGS_{EA}$ and fertility adjusted for different sets of covariates.** Expected fertility ratios (EFR) for the association between $PGS_{EA}$ and number of children in unrelated individuals, shown for the full sample and stratified by sex. All models adjust for year-of-birth categories and the first 10 genetic principal components. Additional covariates are listed in the legend: county of birth (CoB) and wave of participation. The error bars denote 95% confidence intervals. The statistical significance level is denoted by asterisks (* $P < 0.05$, ** $P < 0.01$, *** $P < 0.001$).

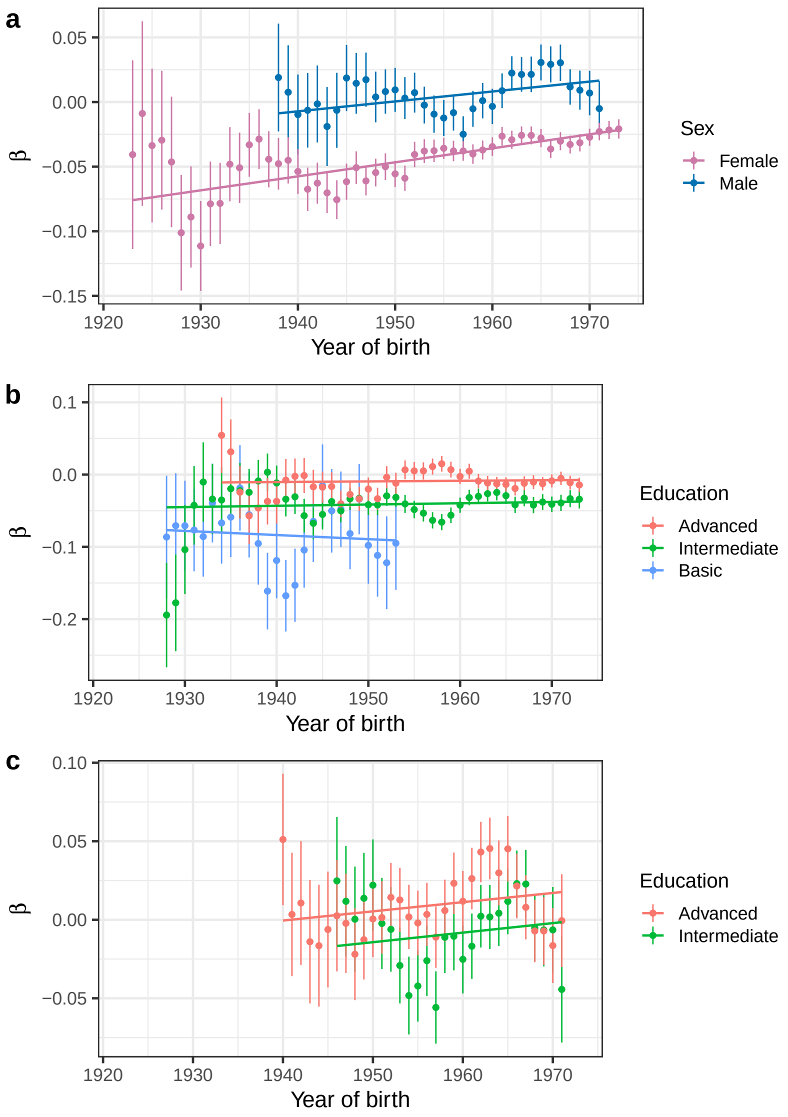


Figure S3: **Effect of $PGS_{EA}$ on completed fertility by year of birth.** Models were estimated (a) by sex using all education categories combined, and stratified by education category among (b) females and (c) males. All regression coefficients were estimated in unrelated individuals. Each estimate corresponds to a 5-year window centered on the respective year; windows including fewer than 200 individuals are not shown. Error bars denote ± SE. Linear trends were assessed using weighted linear regression, with weights inversely proportional to the squared standard errors of $PGS_{EA}$ coefficient estimates. Because the data points are not independent, these trends are shown for visualization purposes only and are not intended for formal statistical inference.

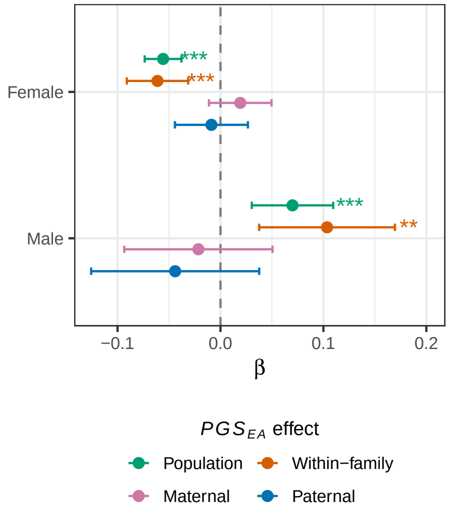


Figure S4: **Within-family genetic effects on fertility estimated using a linear mixed model.** Population-level and within-family effects of individual (proband), maternal, and paternal $PGS_{EA}$ on completed fertility were estimated using snipar. Sibship was treated as a random effect. Models adjusted for year-of-birth categories, county of birth, wave of participation, and the first 10 genetic principal components. Error bars denote 95% confidence intervals. Statistical significance is denoted by asterisks (* $P < 0.05$, ** $P < 0.01$, *** $P < 0.001$).

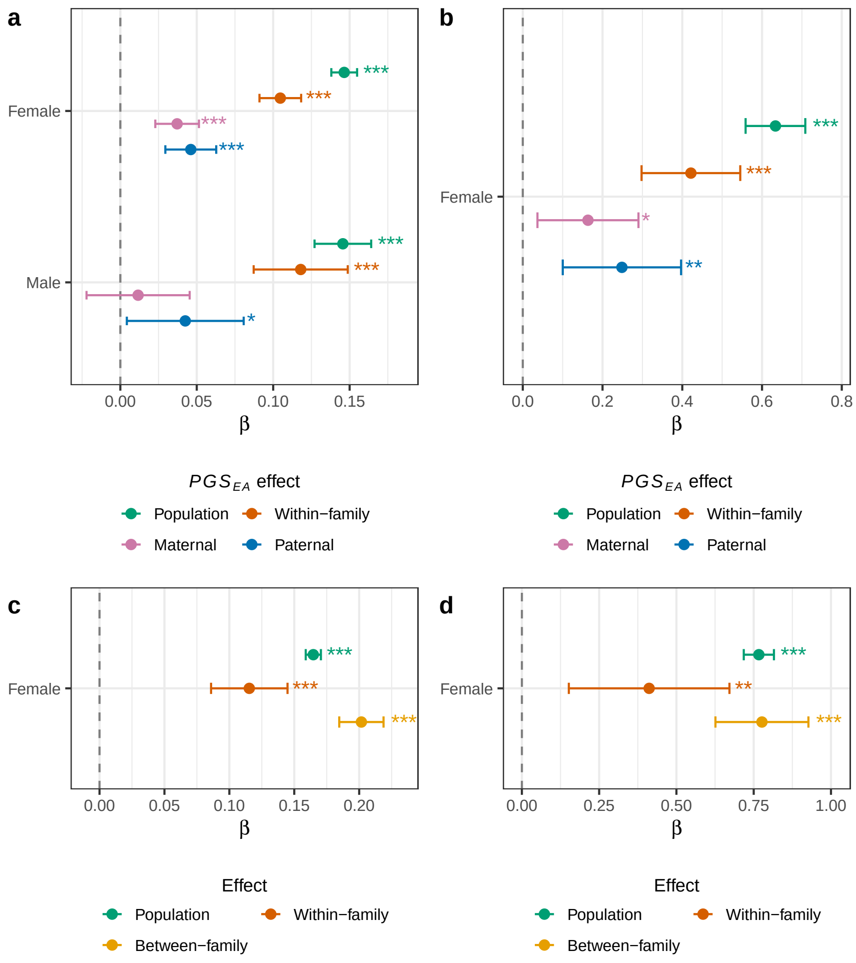


Figure S5: **Within-family genetic effects on educational attainment and age at first pregnancy.** Linear regression coefficients for $PGS_{EA}$ estimated using snipar. Population-level and within-family effects of individual (proband), maternal and paternal $PGS_{EA}$ on (a) EA and (b) AFP. Decomposition of $PGS_{EA}$ effects on (c) EA and (d) AFP in women into population, within-sibling, and between-sibling components. Population models were estimated in unrelated individuals. Models adjust for year-of-birth categories, county of birth, wave of participation, and the first 10 genetic principal components. The error bars denote 95% confidence intervals. The statistical significance level is denoted by asterisks (* $P < 0.05$, ** $P < 0.01$, *** $P < 0.001$).

### Supplementary tables

Table S1: **Educational attainment phenotype definition.** Two definitions of educational attainment were uses: including 11 and 3 categories.

| ISCED 2011 | Complete education category name (11) | EA category (3) | EA value (3) |
|---|---|---|---|
| 0 | Less than primary education | Basic | 0 |
| 1 | Primary education | Basic | 0 |
| 2 | Lower secondary general education | Basic | 0 |
| 2 | Lower secondary vocational education | Basic | 0 |
| 3 | Upper secondary general education | Intermediate | 1 |
| 3 | Upper secondary vocational education | Intermediate | 1 |
| 4 | Post-secondary non-tertiary vocational education | Intermediate | 1 |
| 5 | Short-cycle tertiary education | Intermediate | 1 |
| 6 | Bachelor's or equivalent level | Advanced | 2 |
| 7 | Master's or equivalent level | Advanced | 2 |
| 8 | Doctoral or equivalent level | Advanced | 2 |

Table S2: **Sample size and correlations between polygenic scores, educational attainment (EA) and age at first pregnancy (AFP).** $N$ denotes the sample size with genetic and completed fertility data. Proband $PGS_{EA}$ was available for a subset of individuals for whom parental $PGS_{EA}$ are available. EA phenotype was defined with three possible values: basic = 0; intermediate = 1; advanced = 2.

| Variable | All | Female | Male |
|---|---|---|---|
| $N$ | 46757 | 40738 | 8667 |
| $N_{proband}$ | 11451 | 10214 | 2803 |
| $cor(EA, PGS_{EA})$ | 0.29 | 0.29 | 0.31 |
| $cor(EA, PGS_{EA,proband})$ | 0.27 | 0.27 | 0.28 |
| $cor(EA, PGS_{Cog})$ | 0.16 | 0.16 | 0.17 |
| $cor(EA, PGS_{NonCog})$ | 0.14 | 0.13 | 0.15 |
| $cor(AFP, PGS_{EA})$ | 0.18 | 0.18 | — |
| $cor(AFP, PGS_{EA,proband})$ | 0.15 | 0.16 | — |
| $cor(AFP, PGS_{Cog})$ | 0.07 | 0.07 | — |
| $cor(AFP, PGS_{NonCog})$ | 0.09 | 0.09 | — |
| $cor(PGS_{EA}, PGS_{EA,proband})$ | 0.86 | 0.86 | 0.85 |
| $cor(PGS_{EA}, PGS_{Cog})$ | 0.45 | 0.44 | 0.44 |
| $cor(PGS_{EA}, PGS_{NonCog})$ | 0.49 | 0.49 | 0.50 |
| $cor(PGS_{Cog}, PGS_{NonCog})$ | -0.26 | -0.26 | -0.25 |

Table S3: **Regression results underlying Figure 1c.** Associations between the educational attainment polygenic score ($PGS_{EA}$) and completed fertility, shown for the full sample and stratified by sex, using different sets of covariates. EFRs denote expected fertility ratios; 95% CIs, 95% confidence intervals; YoB, year of birth; 10PCs, top ten genetic principal components; CoB, county of birth; Wave, participation wave.

| Sex | Covariates | EFR | 95% CI | $p$-value | beta | se |
|---|---|---|---|---|---|---|
| All | $YoB + 10PCs$ | 0.968 | $[0.963, 0.973]$ | 3.1e-36 | -0.033 | 0.003 |
| Female | $YoB + 10PCs$ | 0.957 | $[0.951, 0.962]$ | 2.6e-53 | -0.044 | 0.003 |
| Male | $YoB + 10PCs$ | 1.006 | $[0.995, 1.017]$ | 2.9e-01 | 0.006 | 0.006 |
| All | $YoB + 10PCs + CoB$ | 0.972 | $[0.967, 0.977]$ | 4.4e-23 | -0.028 | 0.003 |
| Female | $YoB + 10PCs + CoB$ | 0.961 | $[0.955, 0.967]$ | 4.7e-37 | -0.040 | 0.003 |
| Male | $YoB + 10PCs + CoB$ | 1.007 | $[0.995, 1.02]$ | 2.5e-01 | 0.007 | 0.006 |
| All | $YoB + 10PCs + CoB + Wave$ | 0.974 | $[0.968, 0.979]$ | 8.5e-21 | -0.027 | 0.003 |
| Female | $YoB + 10PCs + CoB + Wave$ | 0.963 | $[0.958, 0.969]$ | 3.8e-33 | -0.037 | 0.003 |
| Male | $YoB + 10PCs + CoB + Wave$ | 1.007 | $[0.995, 1.02]$ | 2.5e-01 | 0.007 | 0.006 |

Table S4: **Interaction between $PGS_{EA}$ and participation wave or historical era.** The table reports results from models testing whether the association between $PGS_{EA}$ and completed fertility varies by wave of participation or historical era. Models were additionally adjusted for EA, and for the interaction term $PGS_{EA} \times EA$ in rows where the EA column is marked "Adj." The interaction term $PGS_{EA} \times Env$ captures heterogeneity in the association across waves or eras. For the era-based analysis, two age thresholds (column "Age") at the time Estonia regained independence in 1991 were used: 18 years (females only) and 25 years (females and males). In the wave-based analysis, a value of 0 corresponds to wave 1 and a value of 1 corresponds to wave 2. In the era-based analysis, a value of 0 corresponds to the Soviet era and a value of 1 corresponds to the post-Soviet era.

| Env | Sex | EA | Age | Predictor | beta | SE | $p$-value |
|---|---|---|---|---|---|---|---|
| Wave | Female | No | – | $PGS_{EA}$ | -0.053 | 0.005 | 1.8e-22 |
| Wave | Female | No | – | Env | -0.072 | 0.007 | 3.5e-22 |
| Wave | Female | No | – | $PGS_{EA} \times Env$ | 0.024 | 0.007 | 3.4e-04 |
| Wave | Male | No | – | $PGS_{EA}$ | 0.029 | 0.014 | 4.3e-02 |
| Wave | Male | No | – | Env | -0.002 | 0.017 | 9.0e-01 |
| Wave | Male | No | – | $PGS_{EA} \times Env$ | -0.027 | 0.016 | 9.2e-02 |
| Wave | Female | Adj. | – | $PGS_{EA}$ | -0.095 | 0.008 | 3.7e-29 |
| Wave | Female | Adj. | – | Env | -0.064 | 0.007 | 9.7e-18 |
| Wave | Female | Adj. | – | $PGS_{EA} \times Env$ | 0.009 | 0.007 | 1.8e-01 |
| Wave | Male | Adj. | – | $PGS_{EA}$ | 0.014 | 0.023 | 5.5e-01 |
| Wave | Male | Adj. | – | Env | -0.003 | 0.017 | 8.7e-01 |
| Wave | Male | Adj. | – | $PGS_{EA} \times Env$ | -0.027 | 0.016 | 8.5e-02 |
| Era | Female | No | 18 | $PGS_{EA}$ | -0.038 | 0.003 | 1.9e-33 |
| Era | Female | No | 18 | Env | -0.169 | 0.121 | 1.6e-01 |
| Era | Female | No | 18 | $PGS_{EA} \times Env$ | 0.023 | 0.014 | 1.1e-01 |
| Era | Female | No | 25 | $PGS_{EA}$ | -0.042 | 0.004 | 1.5e-29 |
| Era | Female | No | 25 | Env | -0.169 | 0.121 | 1.6e-01 |
| Era | Female | No | 25 | $PGS_{EA} \times Env$ | 0.015 | 0.007 | 2.5e-02 |
| Era | Male | No | 25 | $PGS_{EA}$ | 0.006 | 0.007 | 3.9e-01 |
| Era | Male | No | 25 | Env | 0.192 | 0.159 | 2.3e-01 |
| Era | Male | No | 25 | $PGS_{EA} \times Env$ | 0.006 | 0.015 | 7.0e-01 |
| Era | Female | Adj. | 18 | $PGS_{EA}$ | -0.092 | 0.008 | 4.7e-30 |
| Era | Female | Adj. | 18 | Env | -0.107 | 0.120 | 3.7e-01 |
| Era | Female | Adj. | 18 | $PGS_{EA} \times Env$ | 0.014 | 0.014 | 3.2e-01 |
| Era | Female | Adj. | 25 | $PGS_{EA}$ | -0.093 | 0.008 | 5.2e-30 |
| Era | Female | Adj. | 25 | Env | -0.108 | 0.120 | 3.7e-01 |
| Era | Female | Adj. | 25 | $PGS_{EA} \times Env$ | 0.006 | 0.007 | 3.7e-01 |
| Era | Male | Adj. | 25 | $PGS_{EA}$ | -0.011 | 0.019 | 5.8e-01 |
| Era | Male | Adj. | 25 | Env | 0.206 | 0.159 | 1.9e-01 |
| Era | Male | Adj. | 25 | $PGS_{EA} \times Env$ | 0.007 | 0.015 | 6.4e-01 |

Table S5: **Regression results underlying Figure 1d.** Associations between the cognitive and non-cognitive educational attainment polygenic scores ($PGS_{Cog}$ and $PGS_{NonCog}$, respectively) and completed fertility, shown for the full sample and stratified by sex. EFRs denote expected fertility ratios; 95% CIs, 95% confidence intervals.

| Sex | PGS | EFR | 95% CI | $p$-value | beta | se |
|---|---|---|---|---|---|---|
| All | $Cog$ | 0.991 | $[0.986, 0.997]$ | 3.4e-03 | -0.009 | 0.003 |
| Female | $Cog$ | 0.986 | $[0.98, 0.993]$ | 1.6e-05 | -0.014 | 0.003 |
| Male | $Cog$ | 1.007 | $[0.994, 1.019]$ | 3.0e-01 | 0.007 | 0.006 |
| All | $NonCog$ | 0.978 | $[0.973, 0.984]$ | 7.4e-14 | -0.022 | 0.003 |
| Female | $NonCog$ | 0.974 | $[0.968, 0.98]$ | 1.2e-16 | -0.026 | 0.003 |
| Male | $NonCog$ | 0.996 | $[0.984, 1.009]$ | 5.7e-01 | -0.004 | 0.006 |

Table S6: **Regression results underlying Figure 2a.** Associations between the educational attainment polygenic score ($PGS_{EA}$) and completed fertility, estimated in models adjusted for educational attainment, shown for samples stratified by sex. $PGS_{EA}$ *adj.* $EA(3)$ denotes $PGS_{EA}$ adjusted for EA measured using 11 education level categories, whereas $PGS_{EA}$ *adj.* $EA(3)$ denotes $PGS_{EA}$ adjusted for EA measured using 3 categories. EFRs denote expected fertility ratios; 95% CIs, 95% confidence intervals.

| Sex | Predictor | EFR | 95% CI | $p$-value | beta | se |
|---|---|---|---|---|---|---|
| Female | $PGS_{EA}adj.EA(11)$ | 0.974 | $[0.968, 0.98]$ | 9.1e-16 | -0.026 | 0.003 |
| Male | $PGS_{EA}adj.EA(11)$ | 0.998 | $[0.985, 1.011]$ | 7.2e-01 | -0.002 | 0.007 |
| Female | $PGS_{EA}adj.EA(3)$ | 0.974 | $[0.968, 0.98]$ | 6.9e-16 | -0.026 | 0.003 |
| Male | $PGS_{EA}adj.EA(3)$ | 1.001 | $[0.988, 1.014]$ | 8.6e-01 | 0.001 | 0.007 |
| Female | $EA(3)$ | 0.935 | $[0.925, 0.946]$ | 2.1e-31 | -0.067 | 0.006 |
| Male | $EA(3)$ | 1.038 | $[1.013, 1.063]$ | 2.3e-03 | 0.037 | 0.012 |

Table S7: **Regression results underlying Figure 2b.** Associations between the cognitive and non-cognitive educational attainment polygenic scores ($PGS_{Cog}$ and $PGS_{NonCog}$, respectively) and completed fertility, estimated in a model adjusted for educational attainment, shown for samples stratified by sex. EFRs denote expected fertility ratios; 95% CIs, 95% confidence intervals.

| Sex | Predictor | EFR | 95% CI | $p$-value | beta | se |
|---|---|---|---|---|---|---|
| Female | $PGS_{Cog}adj.$ | 0.995 | $[0.988, 1.001]$ | 9.7e-02 | -0.005 | 0.003 |
| Male | $PGS_{Cog}adj.$ | 1.002 | $[0.99, 1.015]$ | 7.3e-01 | 0.002 | 0.007 |
| Female | $PGS_{NonCog}adj.$ | 0.982 | $[0.976, 0.988]$ | 1.0e-08 | -0.018 | 0.003 |
| Male | $PGS_{NonCog}adj.$ | 0.992 | $[0.98, 1.005]$ | 2.4e-01 | -0.008 | 0.007 |

Table S8: **Interaction between $PGS_{EA}$ and educational attainment (EA) or age at first pregnancy (AFP).** Regression results testing whether the association between the educational attainment polygenic score ($PGS_{EA}$) and completed fertility varies by EA or AFP. The interaction term $PGS_{EA} \times Trait$ captures heterogeneity in the association across EA levels or AFP. EA was coded using three categories (basic = 0; intermediate = 1; advanced = 2). AFP was centered within the sample, the mean AFP before centering was 22.6 years.

| Trait | Sex | Predictor | beta | se | $p$-value |
|---|---|---|---|---|---|
| EA | Female | $PGS_{EA}$ | -0.092 | 0.008 | 6.5e-30 |
| EA | Female | $Trait$ | -0.061 | 0.006 | 5.2e-26 |
| EA | Female | $PGS_{EA} \times Trait$ | 0.046 | 0.005 | 7.5e-19 |
| EA | Male | $PGS_{EA}$ | -0.009 | 0.019 | 6.4e-01 |
| EA | Male | $Trait$ | 0.036 | 0.012 | 3.0e-03 |
| EA | Male | $PGS_{EA} \times Trait$ | 0.007 | 0.012 | 5.7e-01 |
| AFP | Female | $PGS_{EA}$ | -0.001 | 0.003 | 7.0e-01 |
| AFP | Female | $Trait$ | -0.031 | 0.001 | 0.0e+00 |
| AFP | Female | $PGS_{EA} \times Trait$ | 0.004 | 0.001 | 2.3e-09 |

Table S9: **Regression results underlying Figure 2c.** Associations between the educational attainment polygenic score ($PGS_{EA}$) and completed fertility, estimated in models stratified by educational attainment, shown for samples stratified by sex. EFRs denote expected fertility ratios; 95% CIs, 95% confidence intervals.

| EA | Sex | EFR | 95% CI | $p$-value | beta | se |
|---|---|---|---|---|---|---|
| All | Female | 0.957 | $[0.951, 0.962]$ | 2.6e-53 | -0.044 | 0.003 |
| Basic | Female | 0.925 | $[0.891, 0.959]$ | 3.2e-05 | -0.078 | 0.019 |
| Intermediate | Female | 0.961 | $[0.952, 0.97]$ | 5.7e-17 | -0.040 | 0.005 |
| Advanced | Female | 0.991 | $[0.983, 1]$ | 4.6e-02 | -0.009 | 0.004 |
| All | Male | 1.006 | $[0.995, 1.017]$ | 2.9e-01 | 0.006 | 0.006 |
| Basic | Male | 1.017 | $[0.873, 1.185]$ | 8.3e-01 | 0.017 | 0.078 |
| Intermediate | Male | 0.992 | $[0.972, 1.012]$ | 4.3e-01 | -0.008 | 0.010 |
| Advanced | Male | 1.008 | $[0.991, 1.026]$ | 3.5e-01 | 0.008 | 0.009 |

Table S10: **Regression results underlying Figure 2d.** Associations between educational attainment (EA) and completed fertility, estimated in models stratified by $PGS_{EA}$, shown for samples stratified by sex. EFRs denote expected fertility ratios; 95% CIs, 95% confidence intervals.

| $PGS_{EA}$ range | Sex | EFR | 95% CI | p-value | beta | se |
|---|---|---|---|---|---|---|
| All | Female | 0.922 | $[0.913, 0.932]$ | 9.5e-49 | -0.081 | 0.005 |
| $(, -1.5)$ | Female | 0.842 | $[0.806, 0.88]$ | 4.2e-14 | -0.172 | 0.023 |
| $[-1.5, -0.5)$ | Female | 0.910 | $[0.889, 0.93]$ | 1.6e-16 | -0.095 | 0.011 |
| $[-0.5, 0.5)$ | Female | 0.944 | $[0.928, 0.961]$ | 1.2e-10 | -0.057 | 0.009 |
| $[0.5, 1.5)$ | Female | 0.968 | $[0.946, 0.991]$ | 7.5e-03 | -0.032 | 0.012 |
| $[1.5, )$ | Female | 0.978 | $[0.923, 1.037]$ | 4.7e-01 | -0.022 | 0.030 |
| All | Male | 1.038 | $[1.015, 1.062]$ | 1.1e-03 | 0.038 | 0.012 |
| $(, -1.5)$ | Male | 1.033 | $[0.896, 1.19]$ | 6.6e-01 | 0.032 | 0.072 |
| $[-1.5, -0.5)$ | Male | 1.017 | $[0.962, 1.075]$ | 5.6e-01 | 0.017 | 0.028 |
| $[-0.5, 0.5)$ | Male | 1.030 | $[0.992, 1.07]$ | 1.2e-01 | 0.030 | 0.019 |
| $[0.5, 1.5)$ | Male | 1.046 | $[1.001, 1.092]$ | 4.7e-02 | 0.045 | 0.022 |
| $[1.5, )$ | Male | 1.092 | $[0.99, 1.203]$ | 7.9e-02 | 0.088 | 0.050 |

Table S11: **Regression results underlying Figure 3a.** Associations between the standard, cognitive and non-cognitive educational attainment polygenic scores ($PGS_{EA}$, $PGS_{Cog}$ and $PGS_{NonCog}$, respectively) and completed fertility among females, estimated in models adjusted for age at first pregnancy (AFP). Besides the standard covariates, the first model included $PGS_{EA}$ and AFP as predictors, while the second model included $PGS_{Cog}$, $PGS_{NonCog}$ and AFP. EFRs denote expected fertility ratios; 95% CIs, 95% confidence intervals.

| Model | Predictor | EFR | 95% CI | *p*-value | beta | se |
|---|---|---|---|---|---|---|
| $PGS_{EA}$ | $PGS_{EA}$ | 0.996 | $[0.991, 1.002]$ | 2.1e-01 | -0.004 | 0.003 |
| $PGS_{EA}$ | $AFP$ | 0.970 | $[0.968, 0.971]$ | <1e-300 | -0.031 | 0.001 |
| $PGS_{Cog/NonCog}$ | $PGS_{Cog}$ | 1.008 | $[1.002, 1.014]$ | 6.2e-03 | 0.008 | 0.003 |
| $PGS_{Cog/NonCog}$ | $PGS_{NonCog}$ | 0.995 | $[0.989, 1]$ | 6.0e-02 | -0.005 | 0.003 |
| $PGS_{Cog/NonCog}$ | $AFP$ | 0.970 | $[0.968, 0.971]$ | <1e-300 | -0.031 | 0.001 |

Table S12: **Results of joint regression model with $PGS_{EA}$, EA, and AFP as predictors.** Associations between $PGS_{EA}$, EA, and AFP and completed fertility among females. EFRs denote expected fertility ratios; 95% CIs, 95% confidence intervals.

| Predictor | EFR | 95% CI | $p$-value | beta | se |
|---|---|---|---|---|---|
| $PGS_{EA}$ | 0.999 | $[0.994, 1.005]$ | 8.4e-01 | -0.001 | 0.003 |
| $EA(3)$ | 0.980 | $[0.970, 0.990]$ | 1.6e-04 | -0.020 | 0.005 |
| $AFP$ | 0.970 | $[0.969, 0.972]$ | $<$1e-300 | -0.030 | 0.001 |

Table S13: **Regression results underlying Figure 3b.** Associations between the educational attainment polygenic score ($PGS_{EA}$) and completed fertility, estimated in models stratified by age at first pregnancy (AFP). Models were estimated with and without additional adjustment for AFP within each stratified group ("AFP adj."). EFRs denote expected fertility ratios; 95% CIs, 95% confidence intervals.

| AFP | Sex | AFP adj. | EFR | 95% CI | $p$-value | beta | se |
|---|---|---|---|---|---|---|---|
| All | Female | False | 0.957 | $[0.951, 0.962]$ | 2.6e-53 | -0.044 | 0.003 |
| With AFP | Female | False | 0.974 | $[0.969, 0.98]$ | 1.9e-19 | -0.026 | 0.003 |
| $(, 20)$ | Female | False | 0.969 | $[0.96, 0.978]$ | 1.1e-10 | -0.032 | 0.005 |
| $[20, 25)$ | Female | False | 1.002 | $[0.994, 1.01]$ | 5.9e-01 | 0.002 | 0.004 |
| $[25, 30)$ | Female | False | 1.020 | $[1.004, 1.036]$ | 1.2e-02 | 0.020 | 0.008 |
| $[30, 35)$ | Female | False | 1.045 | $[1.013, 1.077]$ | 5.3e-03 | 0.044 | 0.016 |
| $[35, )$ | Female | False | 1.006 | $[0.95, 1.066]$ | 8.4e-01 | 0.006 | 0.029 |
| With AFP | Female | True | 0.996 | $[0.991, 1.002]$ | 2.1e-01 | -0.004 | 0.003 |
| $(, 20)$ | Female | True | 0.973 | $[0.963, 0.982]$ | 1.3e-08 | -0.028 | 0.005 |
| $[20, 25)$ | Female | True | 1.005 | $[0.997, 1.013]$ | 2.0e-01 | 0.005 | 0.004 |
| $[25, 30)$ | Female | True | 1.021 | $[1.005, 1.037]$ | 8.8e-03 | 0.021 | 0.008 |
| $[30, 35)$ | Female | True | 1.046 | $[1.014, 1.079]$ | 4.1e-03 | 0.045 | 0.016 |
| $[35, )$ | Female | True | 1.009 | $[0.953, 1.069]$ | 7.6e-01 | 0.009 | 0.029 |

Table S14: **Regression results underlying Figure 3c.** Associations between age at first pregnancy (AFP) and completed fertility among females, estimated in models stratified by $PGS_{EA}$. EFRs denote expected fertility ratios; 95% CIs, 95% confidence intervals.

| $PGS_{EA}$ | Sex | EFR | 95% CI | $p$-value | beta | se |
|---|---|---|---|---|---|---|
| All | Female | 0.970 | $[0.968, 0.971]$ | <1e-300 | -0.031 | 0.001 |
| $(, -1.5)$ | Female | 0.960 | $[0.953, 0.967]$ | 2.8e-28 | -0.041 | 0.004 |
| $[-1.5, -0.5)$ | Female | 0.964 | $[0.961, 0.967]$ | 1.6e-110 | -0.037 | 0.002 |
| $[-0.5, 0.5)$ | Female | 0.972 | $[0.97, 0.974]$ | 2.0e-135 | -0.029 | 0.001 |
| $[0.5, 1.5)$ | Female | 0.973 | $[0.97, 0.975]$ | 3.6e-96 | -0.028 | 0.001 |
| $[1.5, )$ | Female | 0.971 | $[0.966, 0.976]$ | 4.7e-30 | -0.029 | 0.003 |

Table S15: **Regression results underlying Figure 4a.** Associations between the proband (individual's) educational attainment polygenic scores and completed fertility in population and within-family model adjusted for parental PGS, shown for the sample stratified by sex. Fixed-effect quasi-Poisson generalized linear regression models were used. EFRs denote expected fertility ratios; 95% CIs, 95% confidence intervals.

| Sex | Effect | EFR | 95% CI | $p$-value | beta | se |
|---|---|---|---|---|---|---|
| Female | Population | 0.974 | $[0.965, 0.982]$ | 4.9e-10 | -0.027 | 0.004 |
| Female | Direct | 0.971 | $[0.958, 0.986]$ | 8.2e-05 | -0.029 | 0.007 |
| Male | Population | 1.030 | $[1.012, 1.047]$ | 8.0e-04 | 0.029 | 0.009 |
| Male | Direct | 1.043 | $[1.014, 1.074]$ | 4.0e-03 | 0.042 | 0.015 |

Table S16: **Regression results underlying Figure 4b.** Within- and between-family associations between the standard (EA), cognitive (Cog) and non-cognitive (NonCog) educational attainment polygenic scores and completed fertility, estimated in female siblings. EFRs denote expected fertility ratios; 95% CIs, 95% confidence intervals.

| PGS | Effect | EFR | 95% CI | $p$-value | beta | se |
|---|---|---|---|---|---|---|
| EA | Within-family | 0.952 | $[0.924, 0.98]$ | 1.0e-03 | -0.049 | 0.015 |
| EA | Between-family | 0.965 | $[0.949, 0.981]$ | 3.7e-05 | -0.036 | 0.009 |
| Cog | Within-family | 0.985 | $[0.957, 1.014]$ | 3.1e-01 | -0.015 | 0.015 |
| Cog | Between-family | 0.988 | $[0.971, 1.004]$ | 1.5e-01 | -0.013 | 0.009 |
| NonCog | Within-family | 0.974 | $[0.946, 1.004]$ | 8.6e-02 | -0.026 | 0.015 |
| NonCog | Between-family | 0.982 | $[0.965, 0.999]$ | 3.5e-02 | -0.019 | 0.009 |

Table S17: **Regression results underlying Figure S4.** Associations between the proband (individual's), maternal and paternal educational attainment polygenic scores and fertility from snipar model, shown for the sample stratified by sex. EFRs denote expected fertility ratios; 95% CIs, 95% confidence intervals.

| Sex | Effect | beta | se | 95% CI | $p$-value |
|---|---|---|---|---|---|
| Female | Population | -0.056 | 0.009 | $[-0.074, -0.038]$ | 9.7e-10 |
| Female | Within-family | -0.061 | 0.015 | $[-0.091, -0.031]$ | 5.7e-05 |
| Female | Maternal | 0.019 | 0.015 | $[-0.011, 0.050]$ | 2.1e-01 |
| Female | Paternal | -0.009 | 0.018 | $[-0.044, 0.027]$ | 6.3e-01 |
| Male | Population | 0.070 | 0.020 | $[0.030, 0.109]$ | 5.2e-04 |
| Male | Within-family | 0.104 | 0.034 | $[0.038, 0.169]$ | 2.1e-03 |
| Male | Maternal | -0.021 | 0.037 | $[-0.093, 0.051]$ | 5.6e-01 |
| Male | Paternal | -0.044 | 0.042 | $[-0.126, 0.038]$ | 2.9e-01 |

Table S18: **Regression results underlying Figure S5a.** Associations between the proband (individual's), maternal and paternal educational attainment polygenic scores and educational attainment estimated using the snipar model, shown for the sample stratified by sex. EFRs denote expected fertility ratios; 95% CIs, 95% confidence intervals.

| Sex | Effect | beta | se | 95% CI | $p$-value |
|---|---|---|---|---|---|
| Female | Population | 0.146 | 0.004 | $[0.138, 0.155]$ | 4.6e-254 |
| Female | Within-family | 0.105 | 0.007 | $[0.091, 0.118]$ | 3.7e-51 |
| Female | Maternal | 0.037 | 0.007 | $[0.023, 0.051]$ | 3.4e-07 |
| Female | Paternal | 0.046 | 0.008 | $[0.029, 0.063]$ | 5.7e-08 |
| Male | Population | 0.146 | 0.009 | $[0.127, 0.164]$ | 1.7e-53 |
| Male | Within-family | 0.118 | 0.016 | $[0.087, 0.149]$ | 5.6e-14 |
| Male | Maternal | 0.012 | 0.017 | $[-0.022, 0.045]$ | 5.0e-01 |
| Male | Paternal | 0.042 | 0.020 | $[0.004, 0.081]$ | 2.9e-02 |

Table S19: **Regression results underlying Figure S5c.** Within- and between-family associations between the educational attainment polygenic score and educational attainment, estimated in female siblings. EFRs denote expected fertility ratios; 95% CIs, 95% confidence intervals.

| Effect | beta | se | 95% CI | $p$-value |
|---|---|---|---|---|
| Population | 0.165 | 0.003 | $[0.159, 0.171]$ | <1e-300 |
| Within-family | 0.115 | 0.015 | $[0.086, 0.145]$ | 1.9e-14 |
| Between-family | 0.202 | 0.009 | $[0.185, 0.219]$ | 4.6e-113 |

Table S20: **Regression results underlying Figure S5b.** Associations between the proband (individual's), maternal and paternal educational attainment polygenic scores and age at first pregnancy estimated using the snipar model. EFRs denote expected fertility ratios; 95% CIs, 95% confidence intervals.

| Sex | Effect | beta | se | 95% CI | *p*-value |
|---|---|---|---|---|---|
| Female | Population | 0.634 | 0.038 | [0.559, 0.709] | 8.0e-62 |
| Female | Within-family | 0.422 | 0.063 | [0.298, 0.545] | 2.4e-11 |
| Female | Maternal | 0.163 | 0.065 | [0.037, 0.290] | 1.1e-02 |
| Female | Paternal | 0.249 | 0.076 | [0.100, 0.397] | 1.0e-03 |

Table S21: **Regression results underlying Figure S5d.** Within- and between-family associations between the educational attainment polygenic score and age at first pregnancy, estimated in female siblings. EFRs denote expected fertility ratios; 95% CIs, 95% confidence intervals.

| Effect | beta | se | 95% CI | *p*-value |
|---|---|---|---|---|
| Population | 0.767 | 0.025 | [0.718, 0.815] | 1.2e-204 |
| Within-family | 0.412 | 0.133 | [0.152, 0.672] | 1.9e-03 |
| Between-family | 0.777 | 0.077 | [0.626, 0.927] | 8.9e-24 |
